# Age-related loss of brain demyelinating and remyelinating potential is overcome by microglia renewal

**DOI:** 10.64898/2026.06.15.732390

**Authors:** Athena Boutou, Ilias Roufagalas, Irini Papazian, Gianmarco Abbadessa, Konstantina Asprou, Owain W. Howell, Maria Kourouvani, Makoto Sainouchi, Ildiko Farkas, Vasiliki Pogka, Dimitrios Tremoulis, Timokratis Karamitros, Hans Lassmann, Richard Nicholas, Jan Bauer, Lesley Probert

## Abstract

Multiple sclerosis (MS) is an immune-mediated demyelinating disease, with progressive neurodegeneration that is refractory to current therapies. Here, we identify microglia aging as a major factor responsible for age-related loss of beneficial demyelinating and remyelinating brain functions. Brain transcriptomics and in situ spatial gene transcription analysis revealed significant oligodendrocyte loss in both young and aged mice during cuprizone-induced experimental demyelination, but impaired microglial activation in aged mice. Age-related defects in microglial activation were associated with reduced clearance of dead myelin and accumulation of lipid droplets, and impaired remyelination, implying exhaustion of microglial function in the aged mouse brain. Transcriptomic analysis of human brain samples from MS donors with matched disease course and severity and with chronic active and inactive lesions, validated that microglial activation is strongly reduced with increasing age. To investigate microglial responses after repeated demyelinating insults and their direct impact on myelin integrity, we established an experimental model of repeated demyelinating episodes in young and aged mice to recapitulate MS features. Notably, aged mice developed a progressive neuroinflammatory response following sequential demyelinating episodes, in contrast to the alternating cycles of demyelination and remyelination reminiscent of relapsing-remitting MS that were observed in young mice. Microglia depletion and repopulation using a CSF1R antagonist recovered demyelination-remyelination capacity in aged mice. The results indicate that microglia aging is a major determinant in the pathogenesis of progressive MS, and that microglia replacement represents a promising therapeutic approach.

## Introduction

Central nervous system (CNS) myelin is a multilamellar lipid-rich membrane that ensheathes neuronal axons and ensures rapid conduction of electrical signals and metabolic support of neurons^1^. Oligodendrocyte progenitor cells (OPCs) proliferate and differentiate into myelinating oligodendrocytes (OLGs) throughout life as part of myelin maintenance and remodelling^2^. However, aging negatively affects OLGs, myelin integrity and brain white matter (WM) volume^3–5^, as well as spontaneous myelin repair by OPC differentiation and remyelination^6,7^. Brain-wide single cell RNA-sequencing in healthy aged mice revealed increased inflammatory and immune response genes in microglia and astrocytes, and decreased expression of genes involved in cholesterol and lipid metabolism in mature OLGs, further indicating that myelin integrity is compromised with age^8,9^. Age-related changes in myelin are closely linked to cognitive decline and neurological disabilities^4,10,11^, and further contribute to the progression of chronic neurodegenerative disorders such as multiple sclerosis (MS)^12,13^ and Alzheimer’s disease^3,14^.

Microglia are a self-renewing population of CNS macrophages responsible for immune responses and the phagocytic clearance of cellular debris^15^. Healthy microglia are necessary for maintaining the structural and functional integrity of the myelin sheath in adulthood in mice and humans^16,17^. Myelin degradation increases with age, placing an increased burden on microglia to phagocytose myelin debris and initiate remyelination^5^. Spatial and single-nucleus transcriptomics studies of aged mouse brain revealed that myelin-rich WM regions are hotspots for age-related glial cell activation, which in microglia is dependent on their proximity to OLGs^9,18,19^. In demyelination models, microglia accumulate in lesions and are critical for phagocytosing and clearing large amounts of lipid-rich myelin debris that otherwise inhibit remyelination by OPCs^20–24^. Aged microglia show reduced debris clearance and lipid metabolism deficits, leading to poor resolution of neuroinflammation and repair of demyelinated lesions^25–27^. There is significant evidence that microglia dysfunction contributes to the pathogenesis of human MS, including significant numbers of microglial genes associated with MS risk alleles^28^, and the dominance of myelin-containing foamy phagocytes in active, and at the rim of chronic active MS lesions^29,30^. Together, the data support the idea that microglia dysfunction significantly contributes to loss of myelin structure and function in aging and demyelinating diseases.

Myelin debris overloading induces long-term alterations in microglia in both aged and demyelinating brains. In aged mice, microglia accumulate insoluble myelin debris such as lipid droplets, and lipofuscin-like inclusions^5^. Demyelination in aged mice increases debris accumulation leading to the formation of cholesterol-rich crystals, cellular damage and inflammasome activation^26^. Major unanswered questions include whether microglia retain reserve potential after a primary demyelinating event to resolve a second demyelination challenge, how such a response is impacted by age, and whether pharmacologically-induced microglia repopulation might improve this potential. Answers to these questions are relevant for understanding the pathogenesis of human MS, in which myelin pathology is central to the disease. Myelin injury long-precedes disease onset^31,32^, and WM lesions represent pathological substrates for clinical relapses^33^, while disease progression is driven by independent CNS-compartmentalized mechanisms such as chronic neuroinflammation, accumulating axonal damage, and age-related changes in CNS cells^33–35^. In this study we investigated the effects of age on microglial activation and function using *in vitr*o and *in vivo* CNS demyelination models, in particular comparing the functional potential of young and aged microglia to clear and repair myelin following repeated demyelination challenges. We also examined the effects of enhanced microglia turnover in vivo, as a potential therapeutic strategy for limiting chronic neuroinflammatory responses and improving remyelination potential.

## Materials and methods

### MS brain donors and neuropathological assessment

We included 22 UK Brain Bank MS donors (Supplementary Table 1). To minimize confounding by disease course and severity the donors were selected from a total of 167 MS donors through exact matching on time from symptom onset to wheelchair dependence (EDSS 7) and disease duration. All MS donors were treatment-naïve.

Neuropathological assessment was performed on all six sampled CNS regions per donor: superior frontal gyrus GM and WM, cingulate gyrus GM and WM, occipital GM and WM, thalamus at the level of the mammillary bodies, basal pons, and cerebellar WM including the dentate nucleus. Microglia/macrophage density was assessed by HLA-D immunoreactivity using a mouse anti-HLA-DP/DQ/DR antibody (clone CR3/43; Dako, Glostrup, Denmark), whereas myelin integrity was evaluated with an anti-myelin oligodendrocyte glycoprotein antibody (MOG; clone z12, in house). Signal detection was performed with an HRP-conjugated anti-mouse secondary antibody (ImmPRESS HRP, Vector Laboratories, UK) and visualized with diaminobenzidine (ImmPACT DAB, Vector Laboratories, UK). All sections were counterstained with hematoxylin, mounted in DePeX, and digitized at 40-200x using an Aperio AT2 slide scanner (Leica Biosystems, Nussloch, Germany). Lesions were classified as active, chronic active, or chronic inactive. For the present analysis, lesion burden was summarized as the total number of non-cortical lesions and as counts by lesion class. Demyelination was summarized as the mean percent area of all lesions across all sampled regions and separately as percent area of total WM, cortical GM, and deep GM compartments.

## MS brain RNA sequencing

RNA was extracted from pons tissue blocks using the Qiagen RNeasy Lipid Tissue Mini Kit. Libraries were generated with the Lexogen QuantSeq 3′ mRNA FWD kit and sequenced on an Illumina NextSeq 2000 platform. Sequencing data were processed using the nf-core/rnaseq pipeline (v3.3) in Nextflow (v21.04.0); reads were aligned with STAR, sorted with samtools, and quantified at gene level with RSEM to obtain raw counts and transcripts per million (TPM). Pontine microglial activation was quantified from the RNA-seq raw count matrix using AUCell, a rank-based method that estimates the enrichment of a predefined gene set in each sample on the basis of relative expression ordering^36^. The human gene set used for this analysis corresponded to the microglia activation signature derived from the aged versus young naive comparison (**Figure 2Biii; Supplementary Table 2**), and translated to human orthologs using a mouse-to-human mapping approach based on homologene together with org.Mm.eg.db and org.Hs.eg.db R packages. Specifically, the human microglia activation signature used for analysis was TYROBP, ITGAX, CTSS, CTSZ, CTSD, HEXB, TREM2, CLEC7A, C1QA, AXL, APOE, B2M, CCL3, LYZ, P2RY6, LGALS3, IRF8, CX3CR1, CD68 (**Supplementary Table 2**). Group comparisons between younger and older donors were performed using two-sided Wilcoxon rank-sum tests and p<0.05 was considered statistically significant.

**Figure 1.**
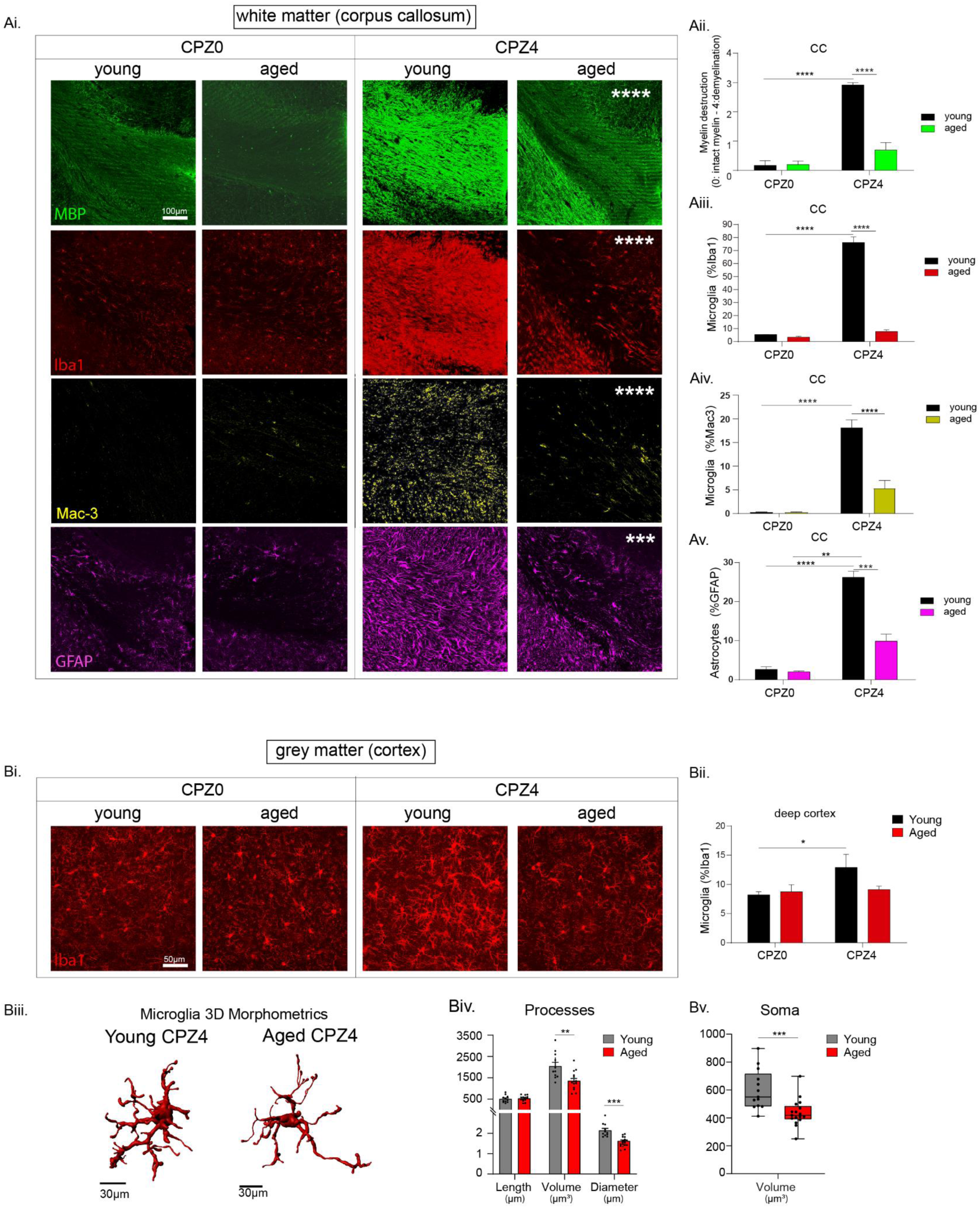
Demyelination and neuroinflammation are impaired in white and grey matter of aged mice **Ai)** Confocal images of midline corpus callosum (CC) white matter in young (2-month-old) and aged (12-month-old) mice at naïve (CPZ0) and demyelination (CPZ4) timepoints showing myelin (MBP-green), microglia (Iba1-red, Mac-3-yellow), and astrocytes (GFAP-magenta). Scale bar: 100μm. **Aii)** Semi-quantitative analysis of myelin integrity score (0: normal myelin – 4: maximum demyelination) in young and aged mice at CPZ0 (naïve) and CPZ4 (demyelination). Quantification of microglia, **Aiii)** as percentage of Iba1-immunoreactivity, **Aiv)** as semi-quantitative score of Mac-3-immunoreactivity, and **Av)** astrocytes as percentage of GFAP-immunoreactivity in young and aged mice at CPZ0 and CPZ4. Bi) Confocal images of cortical grey matter in young (2-month-old) and aged (12-month-old) mice at naïve (CPZ0) and demyelination (CPZ4) timepoints showing microglia (Iba1-red). Scale bar: 50μm. **Bii)** Quantification of cortical microglia as percentage of Iba1-immunoreactivity in young and aged mice at CPZ0 and CPZ4. **Biii)** 3D-cell reconstruction of cortical Iba1^+^ microglia in young (left) and aged (right) at CPZ4. Morphometrics of cortical Iba1^+^ microglia for, **Biv)** cell processes (average diameter: μm, total length: μm, and total volume: μm^3^) and, **Bv)** cell body-soma (volume: μm^3^ and sphericity index) in young and aged mice at CPZ4. (n = 3-5 mice/group, Mean ± SEM, ∗p ≤ 0.05, ∗∗p ≤ 0.001, and ∗∗∗p ≤ 0.0001.)

**Figure 2.**
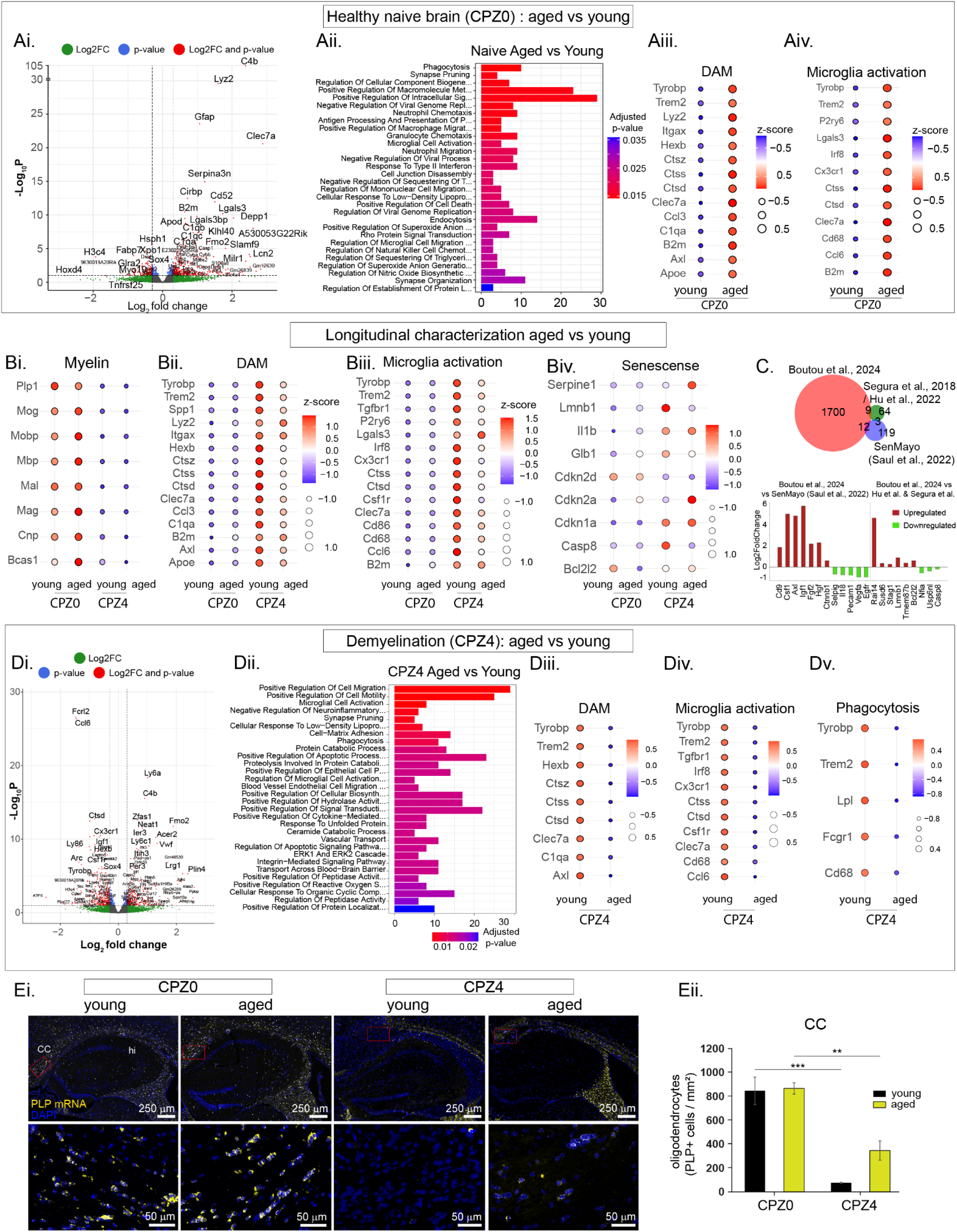
Aged brain transcriptome shows reduced activation responses of primed microglia to cuprizone in the presence of robust oligodendrocyte and myelin damage. Ai-iii) Comparison of RNA sequencing signatures between young and aged healthy naïve brains (CPZ0 aged vs CPZ0 young). **Ai)** Volcano Plot displaying differentially expressed genes (DEGs) in aged vs young naïve (CPZ0) mice. **Aii)** Top regulated statistically significant GO terms for biological processes between aged and young naïve (CPZ0) mice. DotPlots showing z-score for normalized gene counts between young and aged naïve (CPZ0) mice for, **Aiii)** DAM and, **Aiv)** microglia activation statistically significant DEGs. **Bi-iii)** Comparison of brain RNA sequencing signatures between young and aged healthy naïve (CPZ0) and demyelination (CPZ4) brains. DotPlots showing z-score for normalized gene counts between young and aged mice for both naïve (CPZ0) and demyelination (CPZ4) timepoints, for **Bi)** myelin, **Bii)** DAM, **Biii)** microglia activation and **Biv)** cell senescence genes. **C)** Meta-analysis of datasets from our previously published single cell RNA-Seq of cortex from young mice at CPZ0 (naïve) and CPZ3 (demyelination onset) ^23^ with published replicative senescence signatures. From the microglia signature at CPZ3 (1700 DEGs), 9 showed overlap with 64 genes reported by Hernandez-Segura et al., and Hu et al. ^34,35^, and 12 of 119 in the SenMayo cellular senescence gene set reported by Saul et al., ^33^. **Di-v)** Comparison of RNA sequencing signatures between young and aged demyelination brains (CPZ4 aged vs CPZ4 young). **Di)** Volcano Plot displaying differentially expressed genes (DEGs) between young and aged demyelination (CPZ4) groups**. Dii)** Top regulated statistically significant GO terms for biological processes between aged and young CPZ4 mice. DotPlots showing z-score for normalized gene counts between young and aged CPZ4 mice for, **Diii)** DAM, **Div)** microglia activation and, **Dv)** phagocytosis statistically significant DEGs. **Ei)** Spatial distribution of PLP mRNA+ oligodendrocytes by RNAscope in coronal brain sections through corpus callosum (cc) and hippocampus (hi) from young and aged mice at CPZ0 (naïve) and CPZ4 (demyelination) (upper panels), and higher power insets (lower panels) Eii) Quantification of PLP mRNA+ oligodendrocytes/mm3 by RNAscope in the CC of young and aged mice in CPZ0, CPZ4. (n = 5-6 mice/group. ∗p-adjusted ≤ 0.05, ∗∗ p-adjusted ≤ 0.001, and ∗∗∗ p-adjusted ≤ 0.0001).

## Mice

Young (2-month-old), aged (12-month-old) and old (24-month-old) wild-type C57BL/6 (B6) mice were used. Animals were maintained under specific pathogen-free (SPF) conditions at the approved animal facilities of the Department of Animal Models for Biomedical Research, Hellenic Pasteur Institute under the registered codes EL25BIO011, EL25BIO012 and EL25BIO013. All experimental procedures complied to PD 56/2013 and European Directive 2010/63/EU, welfare and ethical use of laboratory animals based on 3+1R: Replacement, Reduction, Refinement and Respect and the guidelines of PREPARE (Planning Research and Experimental Procedures on Animals: Recommendations for Excellence), and ARRIVE (Animal Research: Reporting *in vivo* experiments). The experimental protocol was approved by the Institutional Protocol Evaluation Committee and licensed under the registered codes 266114/28-02-2024 and 71849/30-01-2020 by the National Veterinary Authorities of Attiki Prefecture.

## Experimental model of cuprizone-induced demyelination and remyelination

To enable direct comparisons, young (2-month-old) and aged (12-month-old) male B6 mice were fed *ad libitum* with 0.2% w/w cuprizone (bis-cyclohexanone-oxaldihydrazone; Sigma-Aldrich, catalog C9012) in powdered standard mouse chow for 6 weeks, using a standardized method^23,37,38^. Clinical features of disease were measured in corpus callosum WM and cortical GM at hallmark timepoints determined in the young mice: after 4 weeks (CPZ4, demyelination, acute microgliosis, astrocytosis), 6 weeks (CPZ6, maximum demyelination), and 1 and 4 weeks after cuprizone withdrawal at week 6 (CPZ6+1, early remyelination, CPZ6+4, complete remyelination). An additional chronic protocol of 8 weeks continuous cuprizone feeding (CPZ8) was added for comparisons between young and aged mice. For controls, age-matched healthy untreated brains were used (CPZ0, naïve). In some experiments, to investigate the possibility that reduced biological availability of cuprizone in aged mice might underlie observed differences with young mice, aged mice were fed *ad libitum* with 0.4% w/w cuprizone. For cuprizone experiment 1 (Figure 1), the groups analyzed were CPZ0 controls (young, n=3, aged n=5) and CPZ4 (young, n=3, aged n=5). Group allocations were performed randomly for mice, based on age-matching criteria. Conduction of experiments and IHC analyses were performed by two independent researchers and observations were merged to provide a mean result for the analysis of each experiment.

### “Two-hit” cuprizone-model

A novel chronic “two-hit” cuprizone model was established in mice to induce two consecutive demyelinating episodes separated by a recovery period. Young (2-month-old) and aged (12-month-old) mice were fed with 0.2% cuprizone for 5 weeks (Hit I CPZ5; young n=5, aged n=4) to induce maximum demyelination, allowed to recover without cuprizone for 1 week (Hit I CPZ5+1; young n=4, aged n=5 ), again fed cuprizone for 5 weeks (Hit II CPZ5; young n=5, aged n=6), and again allowed to recover without cuprizone for 1 week (Hit II CPZ5+1; young n=5, aged n=5). Group allocations were performed randomly for mice, based on age-matching criteria. Conduction of experiments and IHC analyses were performed by two independent researchers and observations were merged to provide a mean result for the analysis of each experiment.

## Microglia depletion and repopulation by CSF1R inhibitor

Microglia depletion was induced in vivo using a CSF1R inhibitor (BLZ945, MedChemExpress). BLZ945 induces microglia depletion within 3 days^39^ followed by rapid microglia proliferation and CNS repopulation within 3 days and complete repopulation after 7 days (ref). BLZ945 was administered daily by oral gavage (4 daily doses, 190mg/kg). Young (2-month-old) and aged (12-month-old) mice were fed cuprizone according to the “two-hit” model (above), and treated with BLZ945 starting immediately prior to Hit II CPZ5, to achieve microglia depletion and repopulation prior to the second hit. In this way, we investigated the effects of replenished microglia repopulation upon a second demyelination episode and subsequent remyelination. (Hit II CPZ5; young n=3, aged n=3, Hit II CPZ5+1; young n=2, aged n=3). For untreated controls, animals from the “two-hit” cuprizone model were used for Hit II CPZ5 (young n=5, aged n=6) and Hit II CPZ5+1 (young n=5, aged n=5). The total experimental units n for the BLZ945-treated experiment were n=11. Group allocations were performed randomly for mice, based on age-matching criteria. Experiments and IHC analyses were performed by two independent researchers and observations were merged to provide a mean result for the analysis of each experiment.

## Mouse brain immunofluorescence histochemistry and neuropathology

Mice were transcardially perfused with ice-cold PBS and 4% PFA (pH7.4) followed by overnight post-fixation at 4°C. Brains were dissected and prepared for paraffin, and vibratome sectioning. As previously described^23^, brains were dissected into two hemispheres; the right hemisphere was used for coronal sections to study CC WM pathology (Sidman’s Atlas of the Mouse, sections 295–305), while left hemispheres were used for sagittal sections to study cortical grey matter pathology. For cryostat sectioning, brain hemispheres were cryoprotected by equilibration in ascending sucrose solutions (15% then 30%), freeze-embedded in OCT (Sakura, 4583), and cut into 50 µm serial sections using a cryostat microtome (Leica CM3050S). For vibratome sectioning, 50 µm free-floating serial sections were cut using a vibratome (Leica VT1200). For paraffin sectioning, brains were processed and embedded in paraffin, and 3–5 µm serial coronal sections through the corpus callosum were cut using a microtome (Leica CM3050S). Brain histopathology was performed using primary antibodies, including rat anti-MBP for myelin (1:250, Abcam, ab7349), rabbit anti-Iba1 for microglia (1:1000, Wako chemicals, 019–19741), chicken anti-GFAP for astrocytes (1:1000; Abcam, ab4674). Primary antibodies were incubated overnight at 4°C, followed by fluorescence-labeled secondary antibody including anti-rabbit Alexa Fluor 568 (ThermoFisher Scientific, A-11011), anti-rat Alexa Fluor 647 (Biotium, 20843), anti-chicken Alexa Fluor 488 (Abcam, ab150169) (all 1:1000).

## Multiplex immunofluorescence analysis on paraffin-embedded sections

Multiplex immunofluorescence staining was performed using the Akoya Fluorescent Multiplex kit (Akoya Biosciences) following the manufacturer’s protocol. Briefly, tissue sections underwent antigen retrieval by steaming in pH 6.0 retrieval buffer for 60 min using a household food steamer (Braun), followed by a 10-min blocking step with Opal Antibody Diluent/Block (Akoya Biosciences). Next, the primary antibodies (1:750, Mac-3, BD Biosciences, BDB553322) were incubated for 2 h at room temperature after which sections were washed in Tris-buffered saline with Tween 20 (TBST). Next, horseradish peroxidase (HRP) conjugated donkey a-rat (1:200, Jackson Lab, #712-035-153) was applied for 30 min at room temperature, followed by 10 min. incubation with fluorophore Opal 570. Ultimately the nuclei were stained with 4′,6-diamidino-2-phenylindole (DAPI).

## Quantification of brain immunofluorescence neurohistopathology

Advanced imaging analysis was performed as previously described^23^. Briefly, fluorescent images were acquired as 3D z-stacks (30–35 slices, 1.5 µm steps) were captured from 50 µm sections using 20× and 40× objectives on a Leica TCS SP8 confocal microscope. For tile scans of all cortical layers (I–VI), regions between the corpus callosum and the outer cortex were marked, and 1024×1024 images acquired with a 40× objective were stitched using LAS X software. CC WM was traced using ROI tools in Fiji; in cortical GM, tiles were divided into three regions: upper (I-III), middle (IV-V) and deep (V-VI) cortex. Quantitative analysis was performed using Fiji-ImageJ and IMARIS 9.21.0 (Oxford Instruments). All 3D stacks were resized to ∼30 slices (x = 290.91 µm, y = 290.91 µm, z = 1.5 µm) and converted to 2D maximum-intensity projections (MIPs). Demyelination/remyelination was quantified as MBP-positive area (%) per image volume (mm³) and further assessed using a semi quantitative myelin integrity score based upon hyperfluorescence MBP-positive signal (0, no abnormalities –7, total destruction). Microglia and astrocytes were quantified as Iba1- or GFAP-positive area (%) per image volume (µm³) in MIPs, and as cell density (cells/mm³) by counting positive cells in 3D stacks (Fiji, Cell Counter). Quantification of phagocytosing microglia in the cortical GM was performed by counting the number of cells with evident signal colocalization of Iba1-positive microglia with MBP-positive myelin debris in the 3D z-stack image and is expressed in the figure as the number of phagocytic cells per image volume (μm3).

## Cellular morphometrics

Cellular morphological analysis of microglia was performed semi-automatically, as we have previously described^40^. Individual microglia cells were segmented manually from 3D images acquired by confocal microscopy (Leica SP8), by detailed tracing of the whole cellular body and processes included in each 3D z-stack image, using Fiji/ImageJ2. Cell images were further processed separately in Imaris, and 3D-cell reconstruction was performed using the Surfaces and Filament Tracer plugins. Analysis of 3D reconstruction data was performed automatically as previously described^40^. Cell size quantification was assessed by measurement of cell processes, including total volume (μm^3^), total length (μm) and average diameter (μm). For soma measurements, 3D images from isolated cells were further processed and the cell soma was manually extracted using Fiji’s ROI tool. 3D reconstruction of the cell soma was performed using the Surfaces Plugin in Imaris, with similar specifications applied for each image. IMARIS data measurements for the 3D reconstructed cell soma volume (μm^3^) and sphericity index (index range 0–1, with sphericity index = 1 representing a full sphere), were used for quantification of cell soma size and shape. For 3D image reconstructions of myelin-phagocytosing microglia, dual-channel z-stack images from MBP (myelin) and Iba1 (microglia) were imported in Imaris, split and processed separately using the Surfaces plugin and then merged back together.

## In situ RNAscope hybridization

RNAscope with the PLP mRNA probe was performed according to the Advanced Cell Diagnostics RNAscope Multiplex Fluorescent ReagentKit user manual. In short, paraffin-embedded sections were deparaffinised and treated by a 5-min incubation in target retrieval reagent (ACD, CA) at 95 °C, followed by a 30-min incubation in Protease Plus (ACD, CA) at 40 °C in an oven. Tissue then was incubated with a probe against Proteolipid Protein (*Plp*, 1,057,381-C2, ACD, MA), diluted 1:2000 in TSA buffer for 2 h at 40 °C. This was followed by a three-probe amplification steps (AMP1-AMP3), fluorophore conjugation (Opal 570) and DAPI counterstaining (ACD, CA).

To quantify cells labeled by RNAscope for Proteolipid Protein (PLP) or microglial cells labeled by mac-3, fluorescent stainings were scanned with the Vectra Polaris Automated Quantitative Pathology Imaging System from Perkin Elmer and quantified semi-automatically with QuPath software. To this end, for every slide, the CC WM was traced using ROI tools. In paraffin-embedded sections this was an area of around 0.2 mm2. Next mac-3 immunofluorescence signal was optimized by Qupath’s brightness and contrast tool and quantified as positive area (%) per image volume (mm³). Further, the oligodendrocytes were quantified by manually counting the DAPI+ nuclei with PLP-RNA+ dots in the perinuclear area.

## *In vitro* myelin phagocytosis assays by primary peritoneal macrophages

Myelin phagocytosis assays were performed as previously described^23^. Briefly, primary peritoneal macrophages were harvested from young (2-month-old) and aged (12-month-old) B6 male mice and were incubated with 2.5 μL myelin/well and analyzed after 2, 4, 16, 24h of culture by immunocytochemistry. Cells were fixed on the coverslips with 4% PFA and immunostained using primary antibody markers for myelin (rat anti-MBP, 1:150; Abcam, ab7349), and for macrophages (rabbit anti-Iba1, 1:1000, Wako chemicals, 019–19741) followed by anti-rabbit Alexa Fluor 568 (1:1000 ThermoFisher Scientific, A-11011) anti-rat Alexa Fluor 647 (1:1000, Biotium, 20843) and counterstained with DAPI (Invitrogen). Images were acquired using an SP8 confocal microscope (40x objective, z stack step size 1μm, zoom factor 1.5x). Myelin phagocytosis was measured as the number of Iba1-positive macrophages engulfing MBP-positive myelin debris relative to the total number of Iba1-positive macrophages in the 3D z stack (% of myelin-containing macrophages).

A novel, “two-hit” myelin phagocytosis assay was established to investigate the effects of two consecutive myelin challenges separated by a washout/recovery stage. Primary peritoneal macrophages isolated young (2-month-old) and aged (12-month-old) B6 male mice were incubated with 2.5 μL myelin/well for 16h. Cells were washed with PBS to remove myelin and allow recovery for 24 h washout, and then re-incubated with myelin for a further 4 h, 11 h, and 24 h. Cells were fixed on the coverslips at each time-point during the second myelin phagocytosis challenge, with 4% PFA and immunostained using primary antibody markers for myelin (rat anti-MBP, 1:150; Abcam, ab7349), and for macrophages (rabbit anti-Iba1, 1:1000, Wako chemicals, 019–19741) followed by anti-rabbit Alexa Fluor 568 (1:1000 ThermoFisher Scientific, A-11011) anti-rat Alexa Fluor 647 (1:1000, Biotium, 20843) and counterstained with DAPI (Invitrogen). Images were acquired using an SP8 confocal microscope (40x objective, z stack step size 1μm, zoom factor 1.5x). Myelin phagocytosis was measured as the number of Iba1-positive macrophages engulfing MBP-positive myelin debris relative to the total number of Iba1-positive macrophages in the 3D z stack (% of myelin-containing macrophages).

## Oil Red O for lipid droplets in vitro & in vivo

Lipid droplet accumulation by young and aged macrophages after exposure to myelin in the *in vitro* myelin phagocytosis assay, was measured by Oil Red O (ORO) staining. Primary peritoneal macrophages isolated young (2-month-old) and aged (12-month-old) B6 male mice were incubated with 2.5 μL myelin/well for 16h, followed by myelin washout and incubation with fresh culture medium for further 24h and 48h. Cells were fixed on the coverslips at each time point after myelin exposure with 4% PFA. To verify that both groups had efficient and equal myelin uptake after myelin incubation at 16h, and that myelin was sufficient removed after myelin washout at further 24h and 48h, cells were immunostained for myelin (rat anti-MBP, 1:150; Abcam, ab7349), and for macrophages (rabbit anti-Iba1, 1:1000, Wako chemicals, 019–19741) followed by anti-rabbit Alexa Fluor 568 (1:1000 ThermoFisher Scientific, A-11011) anti-rat Alexa Fluor 647 (1:1000, Biotium, 20843) and counterstained with DAPI (Invitrogen). To measure lipid droplet accumulation, a measure of neutral lipid cargo, cells were stained with ORO at 24h and 48h after myelin wash-out. ORO staining was performed using a freshly prepared ORO working solution and used within 2 h. Briefly, a 0.5% stock solution was prepared by dissolving 0.5 g of Oil Red O powder (Sigma) in 80 ml of 100% isopropanol in a 56 °C water bath overnight, and made up to 100 ml with isopropanol. Prior to use, the stock solution was pre-warmed to 60 °C and filtered. An ORO working solution was prepared by dilution of the stock solution 6:4 with double-distilled water, equilibration for 10 min at room temperature, and filtration with a 0.22 μm membrane (Millipore). Cells were stained with the filtered working solution at 37 °C for 1 min in the dark. ORO accumulation was measured in light microscope images as the number of Iba1-positive macrophages containing ORO+ lipid droplets relative to the total number of Iba1-positive macrophages in the 3D z-stack (%ORO lipid-positive macrophages *in vitro*).

Lipid droplet accumulation was measured by Oil Red O (ORO) staining in frozen sections from in cuprizone-treated young and aged brains for the following groups; CPZ4 (young n = 5, aged n=5), CPZ6 (young n = 5, aged n=5), CPZ6+4 (young n = 5, aged n=5), CPZ8 (young n = 5, aged n=5). Semi-quantitative evaluation of lipid droplet abundance was conducted using a standardized scoring system (0: absence of ORO staining- 3: high ORO lipid accumulation) based on the relative intensity and density of ORO-positive spots observed under a light microscope at 10×,20x magnification. The analysis was carried out separately for two distinct anatomical regions- the corpus callosum (CC) and the adjacent dorsal fornix (FX)- at corresponding rostrocaudal levels (Sidman atlas sections 295–305), allowing the identification of potential region-specific patterns of lipid droplet distribution. Counterstaining with Mayer’s hematoxylin was applied to visualize cell nuclei and facilitate the spatial localization of lipid droplets within the tissue architecture. The resulting semi-quantitative values (ORO+ intensity levels, score 0–3) were used for subsequent graphical and statistical analyses. Group allocations were performed randomly for mice, based on age-matching criteria. Conduction of experiments and IHC analyses were performed by two independent researchers and observations were merged to provide a mean result for the analysis of each experiment.

## Mouse brain RNA-Seq transcriptomics

Brain RNA-Seq was performed in groups of young (2-month-old) and aged (12-month-old) healthy naïve (CPZ0-young, CPZ0-aged) and after 4 weeks of cuprizone-induced demyelination (CPZ4-young, CPZ4-aged), sample size n=6/group. Total brain RNA isolation was performed using TRIzol reagent (Thermo Fisher Scientific, T9424). RNA purity and quantification were assessed using the NanoDrop 2000 spectrophotometer (ThermoFisher Scientific, USA) and the Agilent 2100 Bioanalyzer system (Agilent Technologies, Santa Clara, CA, USA) following the RNA nanochips instructions. Library preparation was performed using the QuantSeq 3′ mRNA-Seq Library Prep Kit-FWD (Lexogen, Vienna, Austria) following the manufacturer’s instructions. Barcode set A was used for indexing, while the number of PCR amplification cycles was determined according to the quantity of each initial RNA sample. Sequencing of the resulting libraries was conducted using the NextSeq2000 platform (Illumina, San Diego, CA, USA) in a 100 bp single-end format. The remaining RNA extracts were stored at −80 °C for future use.

## Bioinformatic analysis for mouse brain RNA-Seq

The read quality of the RNA-seq reads was determined using FastQC v0.12.1. Subsequently, the first 12 bases of the reads were removed and reads with length below 30 were discarded, using bbduk (BBMap v38.94). The reads were aligned to the reference genome GRCm39, using STAR v2.7.9a^41^. Lastly, the reads that correspond to each gene were counted via htseq-count v2.0.3^42^, using the parameters recommended by Lexogen [https://www.lexogen.com/wp-content/uploads/2021/05/015UG108V0311_QuantSeq-Data-Analysis-Pipeline_2021-05-04.pdf].

Differential expression analysis was conducted using DESeq2^43^. Briefly, raw count-matrices and metadata were used as input to DESeq2 function DESeqDataSetFromMatrix to create the DESeq dataset, with a multifactorial design for both age and condition. Pairwise comparisons for all groups was performed using the contrast method. Differential expression results were obtained using the DESeq function with default paameters and differentially expressed (DE) genes were filtered based on the adjusted p-value (≤0.05). Overrepresentation analysis for characterization of enriched GO term gene pathways for biological processes (BP), were retrieved for Mus musculus was performed using clusterProfiler, with GO terms with adjusted p-value <0.05 being deemed significant. All gene expression visualizations were created in R. EnhancedVolcano plot was used for volcano plot visualization of DEG. For dotplots, normalized counts and a calculated z-score per pair comparison were used as input to ggplot2 function for the following signature expressions, filtered by significance (<0.05). Myelin genes *("Mbp", "Mag", "Mog", "Plp1", "Cnp", "Mal", "Bcas1", "Mobp"*), Microglia Activation

> ("Cx3cr1", "Csf1r","Trem2", "Clec7a", "B2m", "Tyrobp", "P2ry6", "Ccl6", "Lgals3", "Irf8",
>
> "Tnf","Ctsd","Ctss","Cd68", "Cd86", "Tgfbr1"), DAM ("Lyz2", "Clec7a","B2m", "Ctss",
>
> "Tyrobp", "C1qa", "Itgax", "Ctsz", "Trem2", "Apoe", "Ccl3", "Ctsd", "Axl", "Hexb"),
>
> senescence ("Cdkn2a", "Cdkn1a", "Cdkn2d", "Casp8", "Il1b", "Glb1", "Serpine1", "Lmnb1", "Bcl2l2").

## Meta-analysis of mouse single cell RNA-Seq with published senescence signatures

Meta-analysis of previously published single cell-RNA- Seq transcriptome datasets for microglia^23^ (GEO Datasets: GSE278137 & GSE278199) was performed by direct comparison of the DEGs of these datasets with recently established senescent gene signatures from previous studies^44,45^ (76 genes),^46^ (134 genes). Genes that were found to overlap with these signatures were visualized using Python with Venn diagrams (venn3 function), and overlapped gene expression direction was visualized in barplots (bar function).

## Statistics

Statistical analyses were performed using Graphpad Prism 8 or MATLAB R 2019b software. Data are presented as mean ± SEM. Optimal sample size was calculated using GPower, based on data from published literature and previous experiments from our labs for measurements for myelin (MBP signal), cellular morphometrics (process diameter, length, volume and soma volume and shape) and bulk RNA-Seq data analyses. For pairwise comparisons Student’s t-test for normally distributed data, or Mann-Whitney for unpaired data were performed. Normality of data was determined using Shapiro-Wilk test. Two-way ANOVA was used for longitudinal analysis or multiple comparisons. For multiple comparisons, *p* values are corrected using the Bonferroni’s or Sidak’s corrections. For transcriptomics bioinformatics analysis FDR-correction for multiple testing was performed using Benjamini and Hochberg method and significance identified using a threshold of FDR ≤0.05. Values of *p* ≤ 0.05 were considered statistically significant (^∗^*p* ≤ 0.05, ^∗∗^*p* ≤ 0.01 ^∗∗∗^*p* ≤ 0.001, ^∗∗∗∗^*p* ≤ 0.0001, ^∗∗∗∗∗^*p* ≤ 0.00001).

## Data availability

Human and mouse RNA-seq data will be deposited in a publicly available database upon publication.

Microscopy data reported in this paper will be shared by the lead contact upon request. This paper does not report original code.

Any additional information required to reanalyze the data reported in this paper will be available from the lead contact upon request.

## Results

### Demyelination and neuroinflammation are impaired in white and grey matter of aged mice

Cuprizone (CPZ)-induced demyelination is a well-characterized and standardized experimental model for studying CNS-compartmentalized mechanisms of neuroinflammation, demyelination and remyelination in mice^38^. It is particularly useful for studying key aspects of progressive MS such as oxidative damage and mitochondrial injury in the presence of chronically activated microglia^47^. As seen by others^48,49^, myelin pathology was significantly impaired in the corpus callosum (CC) WM of aged (12-month-old) compared to young (2-month-old) male C57BL/6 (B6) mice during cuprizone feeding (**Figure 1A**). In young mice, immunofluorescence staining of myelin (MBP) revealed disrupted myelin structure, measured as hyper-intense confluent MBP immunofluorescence by a semi-quantitative myelin destruction score, throughout the midline CC after 4 weeks of cuprizone feeding (CPZ4), compared to young naïve controls (CPZ0) (**Figure 1Ai,ii**). Myelin pathology in young CPZ4 mice was accompanied by highly activated confluent Iba1-positive **(Figure 1Ai,iii)** and Mac-3-positive (**Figure 1Ai,iv**) microglia, as well as reactive GFAP-positive astrocytes (**Figure 1Ai,v**) throughout CC lesions, compared to young CPZ0 mice. While Iba1 immunostaining showed robust microglia activation, the lysosomal marker Mac-3 (LAMP-2, Cd107b) further indicated marked phagocyte activity in young CPZ4 CC. In contrast, in aged CPZ4 mice the integrity of MBP-immunostained myelin in the midline CC appeared relatively unaltered compared to aged CPZ0 mice, and showed a significantly lower destruction score than young CPZ4 mice (**Figure 1Ai,ii**). The reduced myelin pathology was accompanied by less reactive Iba1-positive (**Figure 1Ai,iii**) and Mac-3-positive (**Figure 1Ai,iv**) microglia, as well as GFAP-positive astrocytes (**Figure 1Ai,v**), compared to young CPZ4 mice. Young and aged naïve CPZ0 mice did not show significant differences by MBP-, Iba1-, Mac-3- or GFAP-immunostaining (**Figures 1Ai-iv**).

In the cortical grey matter (GM), no significant alterations in myelin area were detectable in either young or aged mice at CPZ4 (data not shown). Notably, profound demyelination was seen later in young mice, after 5 weeks of cuprizone diet (CPZ5), while aged mice again show little demyelination at this time point (see below **Figure 4C**). Microglia were highly activated in young CPZ4 compared to young CPZ0 mice, but showed no activation in aged CPZ4 compared to aged CPZ0 mice (**Figure 1Bi,ii**). Cell morphometric analysis revealed significant differences between young and aged microglia at CPZ4, with young microglia showing increased process volume and diameter (**Figure 1Biii,iv**), and soma volume (**Figure 1Biii,v**), compared to aged microglia, consistent with increased activation^23,40^. Astrocytes were reactive in both young and aged CPZ4 compared to respective naïve mice, but at much lower levels in aged mice (**Suppl. Figure 1**). The results show that cuprizone-induced myelin pathology affects the CC WM prior to cortical GM in young mice, and is significantly impaired in both regions in aged mice.

**Figure 3.**
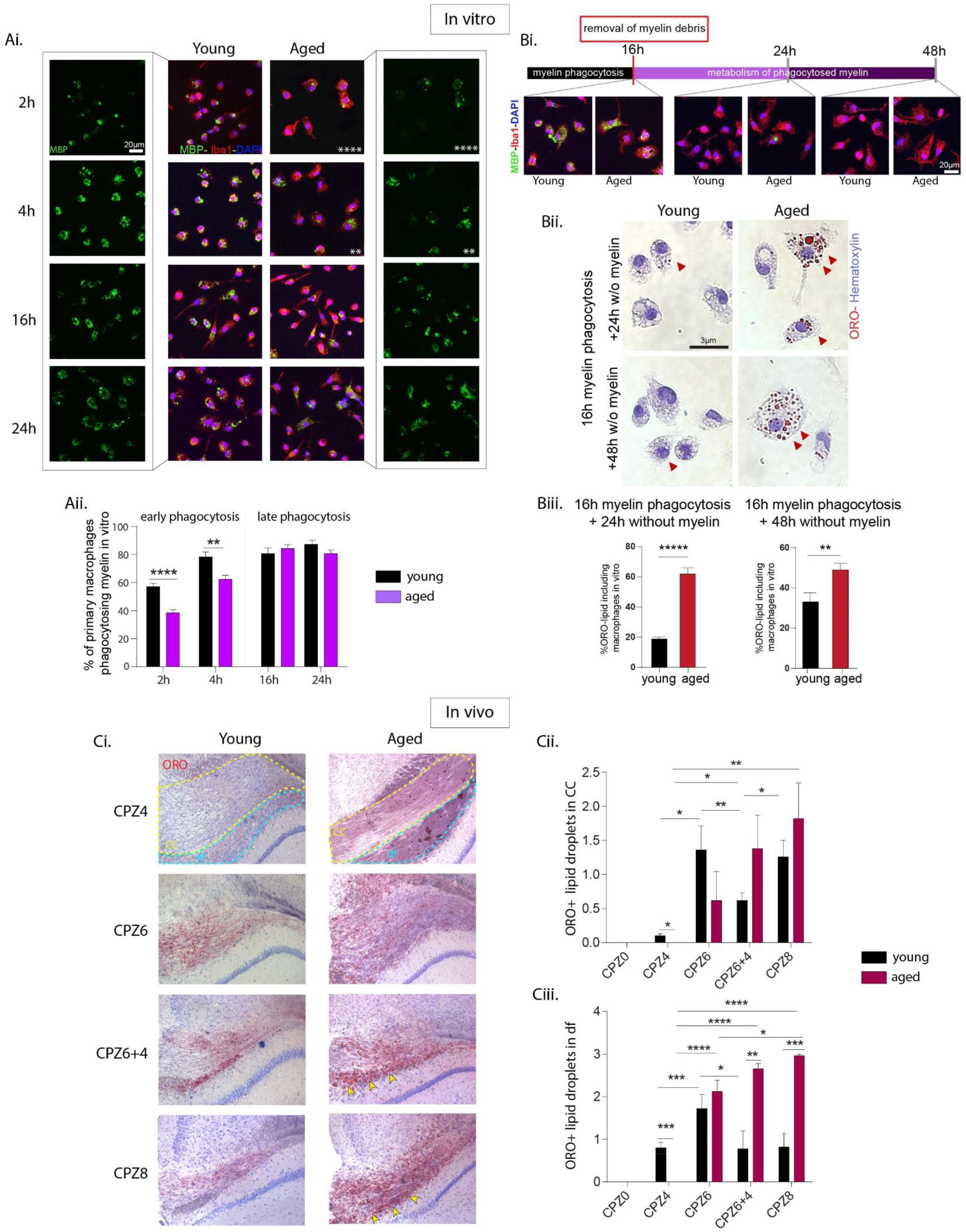
Age-dependent impairment in myelin phagocytosis and lipid droplet clearance in vitro and in vivo **(Ai-ii)** In vitro myelin phagocytosis assay with primary peritoneal macrophages from young (1-month-old) and aged (12-month-old) mice. **(Ai)** Representative immunofluorescence cytochemistry images (myelin basic protein (MBP, green), Iba1 (red), and DAPI (blue) of macrophages from young and aged mice incubated with myelin debris for 2, 4, 16, or 24 h. Scale bar: 20μm. **(Aii)** Quantification of macrophages containing phagocytosed myelin (% of MBP-positive Iba1- macrophages) at early (2–4 h) and late (16–24 h) time points. **(Bi-iii)** Delayed metabolism of phagocytosed myelin by aged macrophages. **(Bi)** Experimental scheme showing removal of myelin from culture medium after 16 h, and assessment of intracellular myelin metabolism by accumulation of Oil-Red-O (ORO) lipid droplets at 24 and 48 h. As an experimental control, representative images of MBP (green), Iba1 (red), and DAPI (blue) (Scale bar: 20μm) in young and aged macrophages showing comparable levels of phagocytosed MBP-immunoreactive myelin in young and aged macrophages at 16 h, and absence of MBP-immunoreactive myelin at 24 h and 48 h, **(Bii)** ORO and hematoxylin staining of young and aged macrophages showing ORO+ lipid droplet accumulation (red, arrowheads). **(Biii)** Quantification of ORO+ lipid–containing macrophages following myelin incubation and withdrawal, demonstrating impaired lipid clearance in aged compared to young macrophages. Scale bar: 3μm. **(Ci-iii)** In vivo accumulation of ORO+ lipid droplets in young (2-month-old) and old (24-month-old) mice during cuprizone-induced demyelination. **(Ci)** Representative ORO-stained coronal brain sections from young and aged mice after 4 (CPZ4), 6 (CPZ6) and 8 (CPZ8) weeks of continuous cuprizone feeding, and CPZ6 followed by 4 weeks of recovery after cuprizone withdrawal (CPZ6+4). Scale bar: 10μm. Quantification of ORO+ lipid droplets in, **(Cii)** the corpus callosum (CC) and **(Ciii)** dorsal fornix (DF) between young and aged mice at CPZ0, CPZ4, CPZ6, CPZ6+4 and CPZ8. (n = 5 mice/group; Mean ± SEM, p<0.05, *p<0.01, **p<0.001, ***p<0.0001).

**Figure 4.**
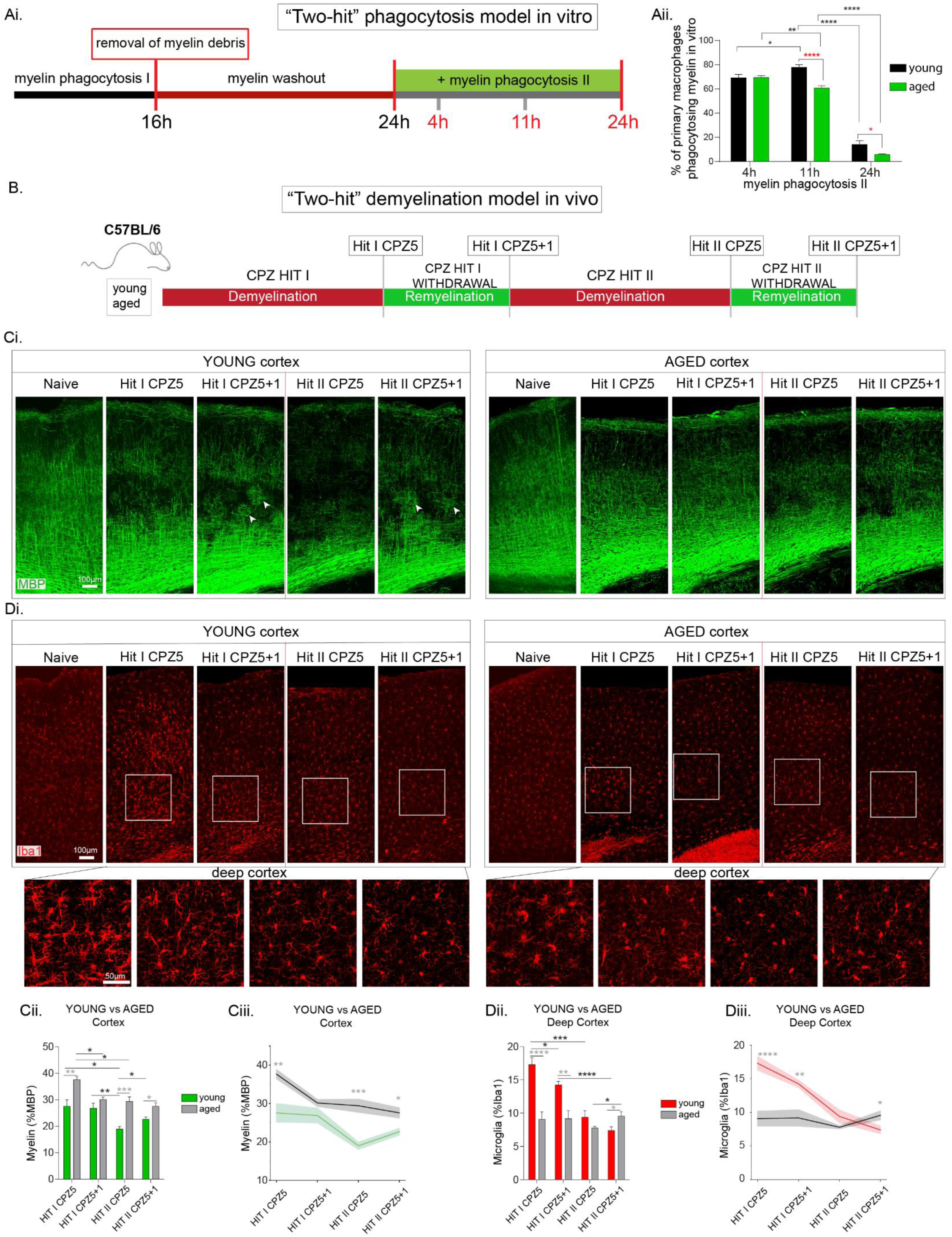
Aged mice show faster phagocyte exhaustion after repeated myelin exposure and failed remyelination in cortical grey matter after sequential demyelination episodes (Ai-ii) Two-hit macrophage myelin phagocytosis model in vitro. **(Ai)** Experimental design: primary peritoneal macrophages incubated with myelin for 16h (Hit I), allowed to recover by removal of extracellular myelin for 24h, and re-incubated with myelin for 4h, 11h, 24h (Hit II). **(Aii)** Quantification of phagocytic capacity of young and aged macrophages measured as MBP-containing, Mac3-immunoreactive cells following the second myelin exposure at 4, 11, and 24h. **(B)** Two-hit cuprizone demyelination model in vivo. Young (2-month-old) and aged (12-month-old) mice were fed 0.2% cuprizone for 5 weeks (Hit I CPZ5) to induce maximum demyelination, allowed to recover without cuprizone for 1 week (Hit I CPZ5+1), again fed cuprizone for 5 weeks (Hit II CPZ5), and allowed to recover for 1 week (Hit II CPZ5+1). **(C-D)** Confocal image-tiles and analysis of myelin and microglia pathology in the cortical grey matter. **(Ci-iii)** Myelin integrity in the cortex during repeated demyelination episodes in young and aged mice. **(Ci)** Representative MBP immunostaining of sagittal sections through somatosensory cortex (layers I-VI) (Scale bar: 100μm) and, **(Cii–Ciii)** quantification of cortical myelin as %MBP immunoreactivity from longitudinal comparison of young vs aged mice following Hit I (CPZ5, CPZ5+1) and Hit II (CPZ5, CPZ5+1). **(Di-iii)** Microglia responses in the cortex during repeated demyelinatiοn episodes in young and aged mice. **(Di)** Representative Iba1 immunostaining in cortical sections from young and aged mice following Hit I (CPZ5, CPZ5+1) and Hit II (CPZ5, CPZ5+1) (Scale bar: 100μm). Insets showing higher magnification images of deep cortical regions (Scale bar: 50μm). **(Dii–Diii)** Quantification of microglia levels (% Iba1 immunoreactivity) from longitudinal comparison of young vs aged mice following Hit I (CPZ5, CPZ5+1) and Hit II (CPZ5, CPZ5+1). (n = 4-6 mice/group; Mean ± SEM, p<0.05, *p<0.01, **p<0.001, ***p<0.0001).

## Aged brain transcriptome reveals reduced activation responses of microglia to cuprizone in the presence of robust OLG and myelin damage

To investigate brain transcriptome changes associated with ageing in cuprizone demyelination, we performed brain RNA-Seq analysis in groups of naïve (CPZ0), and CPZ4 demyelination young (2-month-old) and aged (24-month-old) mice, as well as meta-analysis of previously published single cell RNA-Seq datasets in young mice^23^.

Under naïve conditions, aged CPZ0 brain revealed significant immune activation, and an increased inflammatory gene expression signature compared to young CPZ0 brain (**Figure 2** **A**). Overrepresentation analysis showed that top DEG pathways in aged CPZ0 brain included phagocytosis, synaptic pruning, immune processes and cell death (**Figure 2Aii**). Upregulated genes revealed a prominent signature associated with activated microglia, specifically, genes characteristic of disease-associated microglia (DAM)^50,51^, *(Trem2, Tyrobp, Itgax, Hexb, Clec7a, Axl, Apoe, B2m, Ctss)* in aged CPZ0 compared to young CPZ0 brain (**Figure 2Aiii**). Additionally, genes related to microglia immune activation, including *Cx3cr1, Cd68, Irf8, Ctss, Ctsd, and Ccl6,* were upregulated in aged CPZ0 compared to young CPZ0 brain (**Figure 2Aiv**). Aged CPZ0 brains also showed astrocyte activation, with *Gfap* being among the top upregulated genes (**Figure 2Ai**).

Under demyelination conditions, young and aged brains revealed several marked similarities (**Figure 2Bi-iv**). Both young and aged CPZ4 brains showed prominent downregulation of a myelin/OLG gene signature compared to respective CPZ0 brains, suggestive of significant cuprizone-induced OLG death in aged CPZ4 corpus callosum (**Figure 2Bi**), even though aged brain showed little demyelination by neuropathology at this time point (**Figures 1A**). Both young and aged CPZ4 brains also showed upregulation of microglia activation and DAM gene signatures compared to their respective CPZ0 controls, although at significantly lower levels in aged compared to young brain (**Figure 2Bii-iii**). Both also showed upregulated expression of individual genes associated with cellular senescence compared to respective CPZ0 brains, including *p21/Cdkn1a* in young CPZ4 brain and *p21/Cdkn1a, p16/Cdkn2a* in aged CPZ4 brain, indicative of reduced cell cycling, although neither young nor aged CPZ4 brains showed comprehensive regulation of previously published replicative senescent cell-associated signatures^44–46^ (**Figure 2B**). This was further supported by meta-analysis comparison of our previously published single cell RNA-Seq data at cuprizone demyelination onset in young mice^23^ with published senescence signatures^44–46^ (**Figure 2C**).

Direct comparison between transcriptomes of young and aged CPZ4 demyelination brains revealed a markedly reduced activation response to cuprizone in aged compared to young brains (**Figure 2D**). Consistent with previous reports by our group and others, young CPZ4 brain showed robust upregulation of genes involved in microglia activation, including a DAM signature and phagocytosis^22,23,50–52^, while aged brain showed significantly lower regulation of these gene signatures (**Figure 2D**). Specifically, aged CPZ4 brain showed significantly reduced expression of DAM signature genes *(Trem2, Tyrobp, Hexb, Ctsz, Ctss, Ctsd, Clec7a, C1qa, Axl)* microglia activation *(Tgfbr1, Irf8, Cx3cr1, Csf1r, Ccl6)* and phagocytosis-related genes *(Tyrobp, Lpl, Fcgr1, Cd68)* compared to young CPZ4 brain (**Figure 2Diii-v**).

To validate the RNA-Seq results of prominent downregulation of myelin/OLG genes in response to cuprizone (**Figure 2Bi**), and to interrogate the spatial distribution of transcriptionally-active OLG in the CC of naïve mice, and OLG loss in cuprizone-treated mice, we performed in situ hybridization for PLP mRNA in young and aged brains (**Figure 2E**). Young and aged naïve CPZ0 CC showed equal CC volumes and numbers of PLP mRNA+ OLGs (**Figure 2E**). Both young and aged CPZ4 CC showed increased volumes compared to CPZ0, due to inflammation induced by microglia and astrocytes, and this was more pronounced in young animals. Numbers of PLP mRNA+ OLGs were profoundly reduced in both young and aged CPZ4 CC when volume was taken into account, with loss being significantly greater in young mice (**Figure 2E**).

In short, the transcriptome analysis confirms that microglia in healthy aged brains are already activated (“primed”), but show weaker activation responses to inflammatory stimuli than young microglia. In contrast, myelin and OLGs in young and aged mice are highly susceptible to cuprizone toxicity, even though aged brain showed impaired demyelination by neuropathology. Together, the data are consistent with the idea that reduced microglial responses underlie the impaired initiation of demyelination in aged mice.

## Aged microglia show impaired myelin phagocytosis and lipid clearance

To functionally investigate whether aged microglia show impaired phagocytic clearance of myelin debris, which could underlie the inefficient cuprizone demyelination observed *in vivo* **(Figure 1)**, we set up an *in vitro* myelin phagocytosis model using young and aged macrophages, as previously reported^23^. Peritoneal macrophages were isolated from young (2-month-old) and aged (12-month-old) mice, incubated with myelin debris isolated from wild-type B6 brains for 2 h, 4 h, 16 h, and 24 h, and myelin phagocytic capacity was measured by immunocytochemistry for MBP, Iba1, and DAPI. At early time points (2 h and 4 h), a significant delay in myelin phagocytosis was observed for aged macrophages, measured as a lower percentage of MBP-internalized phagocytic macrophages **(Figure 3Ai,ii).** At later time points (16 h, 24 h), both groups had similar percentages of myelin-containing macrophages **(Figure 3Ai,ii).** This finding aligns with the *in vivo* observation of delayed microglial activation in the aged brain at CPZ4 **(Figure 1)**.

Aged microglia display impaired lipid metabolism^5,25–27^. To investigate whether reduced myelin phagocytosis in aged macrophages is accompanied by alterations in lipid metabolism, we included a myelin washout step in the *in vitro* myelin phagocytosis assay, and measured lipid clearance from the cells. Peritoneal macrophages isolated from young and aged mice were incubated with myelin debris for 16 h, myelin was removed, and the cells were incubated without myelin for 24 h or 48 h and stained with Oil Red O (ORO) to detect lipid droplets. Double immunofluorescence staining for Iba1 and MBP verified that both groups had efficient and equal myelin uptake at 16 h, and lacked MBP immunoreactivity after myelin washout **(Figure 3Bi).** Aged mice, however, showed significantly higher levels of ORO+ lipid-containing macrophages, indicating impaired lipid turnover, compared to young mice both 24 h and 48 h after myelin removal **(Figure 3Bii,iii).**

To investigate the *in vivo* relevance of these findings for brain microglia, we measured lipid droplet accumulation by ORO staining in the CC of young (2-month-old) and aged (12-month-old) mice at naïve (CPZ0), and cuprizone disease time points at 4 (CPZ4), 6 (CPZ6), and 8 (CPZ8) weeks of continued cuprizone feeding, and at 4 weeks after cuprizone withdrawal (CPZ6+4). In young mice, ORO+ signal was absent at naïve (CPZ0), showed few cells with lipid droplets at CPZ4 (peak demyelination in young mice), and robust accumulation of ORO+ cells at CPZ6 and CPZ8 of continuous cuprizone feeding (**Figure 3Ci,ii**). ORO+ signal was significantly reduced when cuprizone was withdrawn at 6 weeks followed by 4 weeks recovery (CPZ6+4) (**Figure 3Ci,ii**), a time point associated with almost complete remyelination in this model^23^. In contrast, in aged mice ORO+ lipid droplets were scarce at CPZ0 and CPZ4, but significantly increased at CPZ6 (onset of demyelination in aged mice) and CPZ8 (demyelination peak in aged mice) of continuous cuprizone feeding, and continued to accumulate after cuprizone withdrawal (CPZ6+4) and in sentinel mice up to 10 weeks after cuprizone withdrawal (data not shown). Notably in aged mice, a particularly high ORO+ signal was observed in the dorsal fornix (df) (**Figure 3Ci, arrows, Ciii**). The density of ORO+ lipid-laden cells in the mouse brain increases with natural aging^53^, and here we observed that numbers dramatically increased and are poorly resolved in aged WM following a demyelination event, reminiscent of accumulations of foamy macrophages/microglia in active and chronic active MS lesions^29,30^. The results show that aged macrophages *in vitro* and microglia *in vivo* phagocytose myelin poorly and accumulate more lipid droplets compared to young phagocytes, a finding consistent with their reduced functional capacity.

## Aged-related impaired microglial responses and failed remyelination between repeated demyelination episodes

In early stages of human MS, demyelinated lesions can undergo spontaneous repair by remyelination, but this process eventually fails^12,13^. To investigate the effects of age on the reserve potential of microglia to phagocytose and clear myelin debris between repeated episodes of demyelination, we introduced a second myelin incubation to the *in vitro* phagocytosis assay. Peritoneal macrophages were isolated from young (2-month-old) and aged (12-month-old) mice, incubated with myelin for 16 h, allowed to recover for a 24 h washout, and re-incubated with myelin for 4 h, 11 h, and 24 h (“two-hit” myelin phagocytosis model) **(Figure 4Ai).** Young macrophages showed increased intracellular MBP-immunoreactivity between 4 and 16 h of the second myelin challenge, while aged macrophages, which already contained significantly more ORO+ lipid droplets than young macrophages after the first myelin challenge (**Figure 3**), showed progressively reduced myelin phagocytic potential during the entire second myelin challenge **(Figure 4Aii)**. Notably, both young and aged macrophages exhibited markedly reduced myelin phagocytosis potential 24 h into the second myelin challenge **(Figure 4Aii)** compared to the first (**Figure 3A**), indicative of phagocytic exhaustion upon repeated exposure to myelin debris *in vitro*.

To investigate the effects of age on the functional capacity of microglia to respond and repair repeated episodes of demyelination *in vivo*, we established a chronic “two-hit” cuprizone demyelination model in mice. Young (2-month-old) and aged (12-month-old) mice were fed cuprizone for 5 weeks (Hit I CPZ5), allowed to recover without cuprizone for 1 week (Hit I CPZ5+1), again fed cuprizone for 5 weeks (Hit II CPZ5), and again allowed to recover without cuprizone for 1 week (Hit II CPZ5+1) (**Figure 4B**). Demyelination and remyelination (MBP), microglia activation (Iba1) and astrocyte reactivity (GFAP) were monitored at each time point in sagittal slices through the somatosensory cortex (**Figure 4**), and coronal slices through the CC (**Figure 5**).

**Figure 5.**
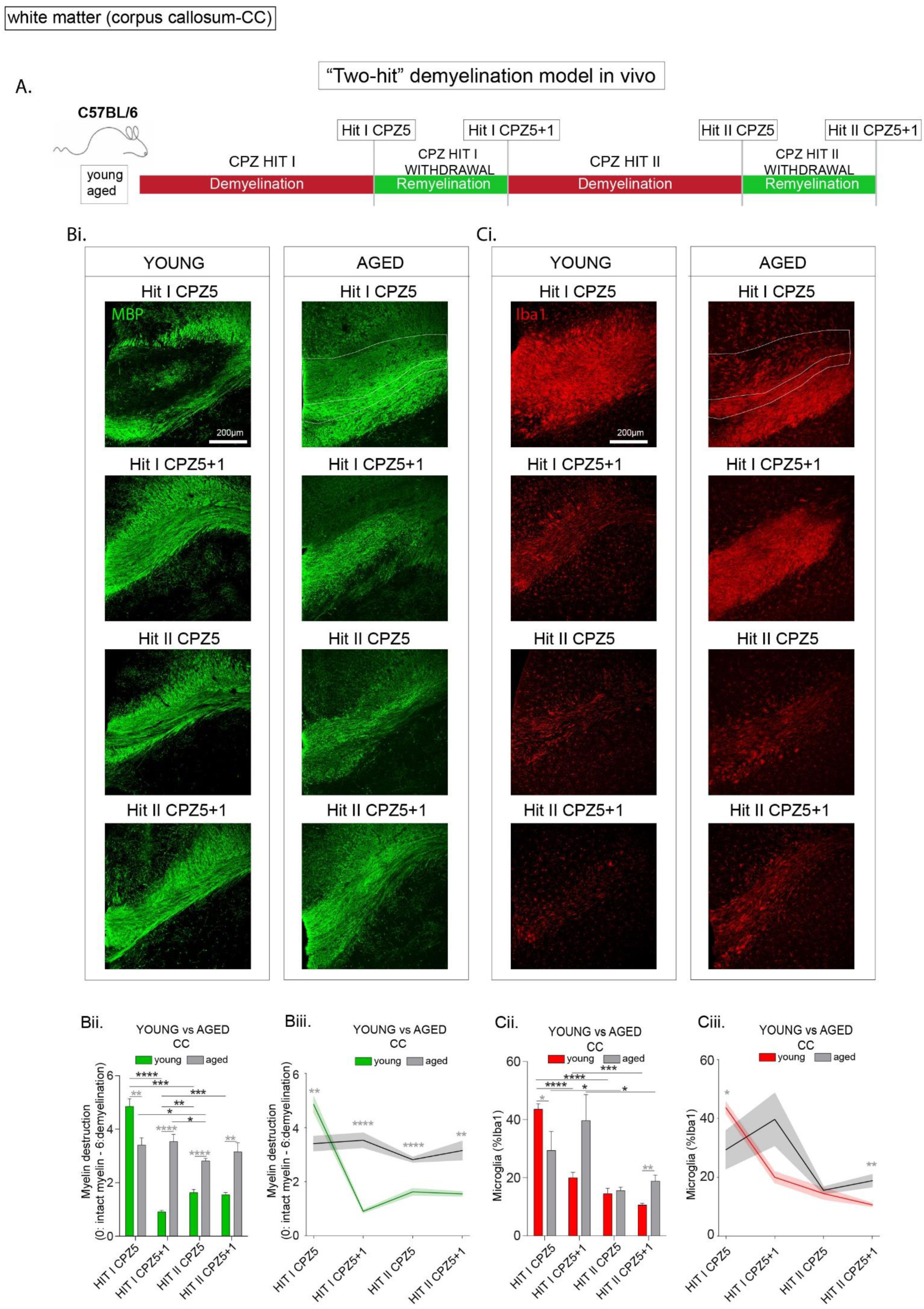
Aged mice show impaired demyelination, remyelination and microglia responses in corpus callosum white matter after repeated demyelination episodes **(A)** Schematic of the two-hit cuprizone demyelination model in vivo. Young (2-month-old) and aged (12-month-old) mice were fed 0.2% cuprizone for 5 weeks (Hit I CPZ5) to induce maximum demyelination, allowed to recover without cuprizone for 1 week (Hit I CPZ5+1), again fed cuprizone for 5 weeks (Hit II CPZ5), and again allowed to recover for 1 week (Hit II CPZ5+1). **(Bi-iii)** Myelin integrity in midline corpus callosum (CC) during repeated demyelinating episodes in young and aged mice. **(Bi)** Representative MBP immunostaining of coronal CC sections in young and aged mice (Scale bar: 200μm) and, **(Bii–Biii)** quantification of demyelination (semi-quantitative score 0: normal intact myelin 6: total myelin disruption) from longitudinal comparison of young vs aged mice following Hit I (CPZ5, CPZ5+1) and Hit II (CPZ5, CPZ5+1). **(Ci-iii)** Microglial response in midline CC during repeated demyelinating episodes in young and aged mice. **(Ci)** Representative Iba1 immunostaining of coronal CC sections from young and aged mice following Hit I (CPZ5, CPZ5+1) and Hit II (CPZ5, CPZ5+1) (Scale bar: 200μm). **(Cii–Ciii)** Quantification of microglia levels (% Iba1 immunoreactivity) from longitudinal comparison of young vs aged mice following Hit I (CPZ5, CPZ5+1) and Hit II (CPZ5, CPZ5+1). (n = 4-6 mice/group; mean ± SEM, p<0.05, *p<0.01, **p<0.001, ***p<0.0001).

In cortical GM, young mice showed efficient demyelination after each cuprizone challenge (**Figure 4C; Hits I&II CPZ5**), followed by significant remyelination after the second demyelination episode (**Figure 4Ci,iii; Hits I&II CPZ5+1, arrowheads)**. Cortical demyelination was greater after the second demyelination episode, compared to the first (**Figure 4Ci,ii)**. In young mice, demyelination was associated with a robust microglia activation response to the first cuprizone challenge (**Figure 4D; Hit I CPZ5**), followed, by gradual reduction of the microglial response, which continued even through the second active demyelination episode (**Figure 4Di,iii)**. Astrocyte reactivity also peaked in response to the first cuprizone hit, subsequently resolved and remained stable thereafter (**Supplementary Fig. 1Bi-iii**). In contrast, aged mice developed a chronic low-level demyelinating disease (**Figure 4Ci-iii; Hits I&II CPZ5**), in which initial demyelination was less than in young mice and continued through the recovery periods after cuprizone withdrawal, with no detectable remyelination activity (**Figure 4Ci-iii; Hits I&II CPZ5+1**). The progressive, low-grade demyelination in aged mice was associated with poor cortical microglia activation and resolution responses throughout the entire disease and recovery course (**Figure 4Di-iii**). Astrocyte reactivity was initially low and tended to gradually increase over subsequent time points (**Supplementary Fig. 1Bi-iii**).

In CC WM, young mice showed characteristic robust demyelination followed by almost complete remyelination after the first cuprizone challenge (**Figure 5Bi-ii; Hit I CPZ5 & CPZ5+1**). Demyelination and remyelination also occurred, albeit at lower levels, after the second cuprizone challenge (**Figure 5Bi-iii; Hit II CPZ5 & CPZ5+1).** As in cortex, demyelination in young CC was associated with a robust microglial activation response to the first cuprizone challenge (**Figure 5Ci-iii; Hit I CPZ5**) followed by gradual reduction of the microglial response, which continued through the second demyelination challenge (**Figure 5Ci-iii)**. Astrocyte reactivity was at equal levels at all time points (**Supplementary Fig. 1Ci-iii**). In contrast, aged mice developed a low-grade progressive demyelinating disease throughout the entire experimental protocol (**Figure 5Bi-iii; Hits I&II CPZ5**). Initial myelin damage was less than in young mice, and persisted through the first recovery period into the second cuprizone challenge and recovery period, with no detectable remyelination activity in the time course of this experiment (**Figure 5Bi-iii; Hits I&II CPZ5+1**). Microglial activation was weak after the first cuprizone challenge, increased in the first recovery period and subsequently reduced and stayed at low levels throughout the second cuprizone challenge and recovery period (**Figure 5Bi-iii**). As in aged cortical GM, astrocyte reactivity was initially low and tended to gradually increase over subsequent time points (**Supplementary Fig. 1Ci-iii**).

Notably in young mice, the pattern of pathology was different in the CC WM and cortical GM, with CC showing maximal neuroinflammation and demyelination after the first cuprizone hit (**Figure 5**) while cortex showed maximum demyelination after the second hit (**Figure 4**), possibly reflecting the increased susceptibility of WM to immune-mediated damage in mice and humans^9,18,19,54^, as well as known heterogeneity of microglia between different brain regions^55^. Notable also is the finding that microglia responses to the second cuprizone challenge were weak compared to the first challenge in both young cortex and CC, suggestive of functional exhaustion (**Figures 4** **& 5**). Overall, the results show that aged mice mount defective microglial responses, characterized by weak activation and resolution potential, which results in impaired demyelination, and subsequently failed remyelination, between repeated demyelination episodes.

## Microglia repopulation after repeated episodes of demyelination promotes remyelination in young and aged mice

To investigate the therapeutic potential of microglial repopulation in remyelination after two independent episodes of demyelination in young and aged mice, we depleted microglia *in vivo* by administration of a CSF1R inhibitor. Young (2-month-old) and aged (12-month-old) mice were fed cuprizone for 5 weeks to induce demyelination (Hit I CPZ5), followed by 1 week of cuprizone withdrawal to allow recovery (Hit I CPZ5+1). Subsequently, animals were exposed to a second 5-week cuprizone challenge (Hit II CPZ5), followed by a second 1 week of recovery (Hit II CPZ5+1). Acute microglia depletion and repopulation was induced using BLZ945, a CSF1R inhibitor^39^, before the second cuprizone hit, to assess the contribution of newly-divided microglia to demyelination and remyelination potential (**Figure 6A)**.

**Figure 6.**
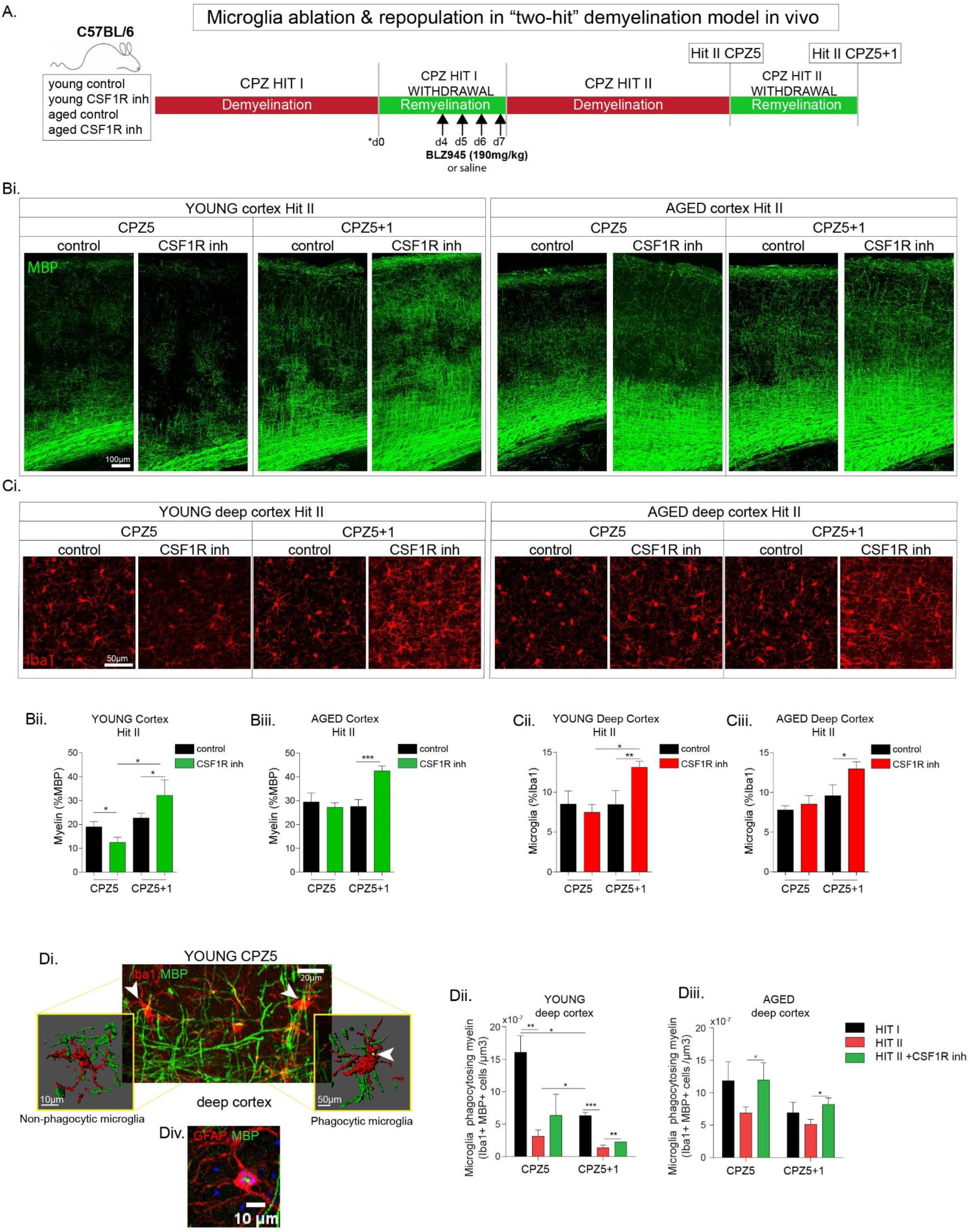
Microglia repopulation following repeated episodes of demyelination promotes remyelination in both young and aged mice **A)** Experimental design of microglia ablation with BLZ945, and repopulation prior to a second cycle of demyelination and remyelination. Young (2-month-old) and aged (12-month-old) mice were fed 0.2% cuprizone for 5 weeks to induce demyelination (Hit I CPZ5), allowed to recover without cuprizone for 1 week (Hit I CPZ5+1), again fed cuprizone for 5 weeks (Hit II CPZ5), and allowed to recover for 1 week (Hit II CPZ5+1). Acute microglia depletion and repopulation was induced by administration of BLZ945, a CSF1R inhibitor, before the second cuprizone hit. Controls were injected with saline vehicle. d0, days of cuprizone withdrawal/ remyelination. **(B-C)** Confocal image-tiles and analysis of myelin and microglia pathology in the cortical grey matter. **(Bi-ii)** Myelin integrity in the cortex during repeated demyelinating episodes and microglia repopulation in young and aged mice. **(Bi)** Representative MBP immunostaining of sagittal sections through somatosensory cortex (layers I-VI) (Scale bar: 100μm) and, quantification of cortical myelin as %MBP immunoreactivity from **(Bii)** young, **(Biii)** aged, at Hit II (CPZ5, CPZ5+1), following treatment with CSF1R inhibitor or saline control. **(Ci-ii)** Microglia responses in the cortex after microglia repopulation during repeated demyelinating episodes in young and aged mice. **(Ci)** Representative Iba1 immunostaining of sagittal sections through somatosensory deep cortex (layers V-VI) (Scale bar: 50μm) and **(Cii–iii)** quantification of microglia as %Iba1-immunoreactivity from **(Cii)** young and**(Ciii)** aged mice at Hit II (CPZ5, CPZ5+1), following treatment with CSF1R inhibitor or saline control. **(Di-iii)** Myelin phagocytosis by microglia of young vs aged mice following Hit I (CPZ5, CPZ5+1) and Hit II (CPZ5, CPZ5+1) and treated with CSF1R inhibitor or saline control in vivo. **(Di)** Representative images (Scale bar: 20μm) and 3D reconstruction (Scale bar: 50μm) of non-phagocytic (left) and myelin-phagocytic microglia (right) depicted as MBP (green)-containing phagocytic Iba1-positive (red) cell and measured in **(Dii)** young and **(Diii)** aged brains as the cell number of MBP+ containing Iba1+ microglia/μm^3^ following Hit I (CPZ5, CPZ5+1) and Hit II (CPZ5, CPZ5+1) and treated with CSF1R inhibitor or saline control **(Div).** Characteristic in vivo image of a GFAP-positive (red) astrocyte containing MBP-positive myelin debris (green) in the cortex at Hit II CPZ5 (Scale bar: 10μm). (n = 2-3 mice/group; mean ± SEM, p<0.05, *p<0.01, **p<0.001, ***p<0.0001).

In the cortical GM of young mice, microglial depletion and repopulation increased demyelination after the second cuprizone hit (**Figure 6Bi,ii; Hit II, CPZ5**), and increased myelin recovery after the second cuprizone withdrawal (**Figure 6Bi,ii; Hit II, CPZ5+1).** Remyelination was associated with significantly increased microglia activation (**Figure 6Ci,ii; Hit II, CPZ5+1).** Notably in the young cortex, microglia repopulation increased the myelin phagocytosis activity of microglia both after the second cuprizone hit (**Figure 6Di-ii**), and during the remyelination phase after the second cuprizone withdrawal (**Figure 6Dii**). Sparse GFAP-positive astrocytes containing MBP-positive myelin were also observed in the cortex of young mice after the second cuprizone challenge (**Figure 6Div**). Similarly, in aged mice, although microglia depletion and repopulation did not alter demyelination after the second cuprizone hit (**Figure 6Bi,iii; Hit II, CPZ5**), it dramatically increased myelin recovery after the second cuprizone withdrawal (**Figure 6Bi,iii; Hit II, CPZ5+1).** As in the young cortex, remyelination was associated with significantly increased microglia activation (**Figure 6i,iii; Hit II, CPZ5+1).** Also, microglia repopulation in aged mice increased the myelin phagocytosis activity of microglia both after the second cuprizone hit (**Figure 6Diii; Hit II, CPZ5**), and during the remyelination phase after the second cuprizone withdrawal (**Figure 6Diii; CPZ5+1**), to levels equal to those seen in the first hit. Together, these results show that microglia depletion by CSF1R inhibition and repopulation by endogenous cells is sufficient to restore functional microglia activity in demyelination and remyelination in both young and aged mice.

## Human MS brain microglia activation is reduced with increasing age

To investigate whether the reduction in microglial activity observed in aged compared to young cuprizone-treated mice is also evident in human MS, we performed RNA-Seq in pons from human MS donors from a cohort of 22 UK Brain Bank MS donors (**Figure 7A, Supplementary Table 1**). Donors were selected from a larger cohort by exact matching for disease duration and time to EDSS 7 to control for disease severity (see Materials and Methods). Lesions were classified based on HLA-D immunoreactivity into active, chronic active (including broad-rim lesions), and chronic inactive, enabling region-specific assessment of microglia/macrophage density and demyelination (**Figure 7B, C**). Subsequently, RNA sequencing of pontine tissue was performed, and microglial activation was quantified using a gene signature derived from the aged versus young mouse cuprizone demyelination comparison (**Fig2Biii, Supplementary Table 2)**, allowing transcriptomic evaluation of age-associated differences in microglial activity. In the age-matched comparison, younger donors presented with more (total) lesions than older donors during the same study period(s) between symptom onset to wheelchair dependence (EDSS 7, median 12 years), and onset to autopsy sampling (median 26 years) (**Figure 7D, Supplementary Table 1**). This difference was mainly driven by chronic active lesions, which were more frequent in the younger group (median 7 vs 2, p=0.0057), together with a higher overall lesion count (median 10 vs 4, p=0.0061) (**Figure 7D, F**). Active lesion counts (**Figure 7E**), and cortical and non-cortical grey matter demyelination (**Figure 7I, J**) showed similar directional trends but did not reach significance (p=0.0538, p=0.13 and p=0.0568, respectively). No clear group difference was observed for inactive lesion counts (**Figure 7G**) or WM demyelination (**Figure 7H**). Pontine microglial activation, quantified by HLA-D immunohistochemistry (**Figure 7C**) and AUCell enrichment of the microglia activation signature (**Figure 7K**), was higher in younger cases (median 0.129 vs 0.081, p=0.0302). Overall, younger MS donors were characterized by a more inflammatory lesion profile and increased pontine microglial activation.

**Figure 7.**
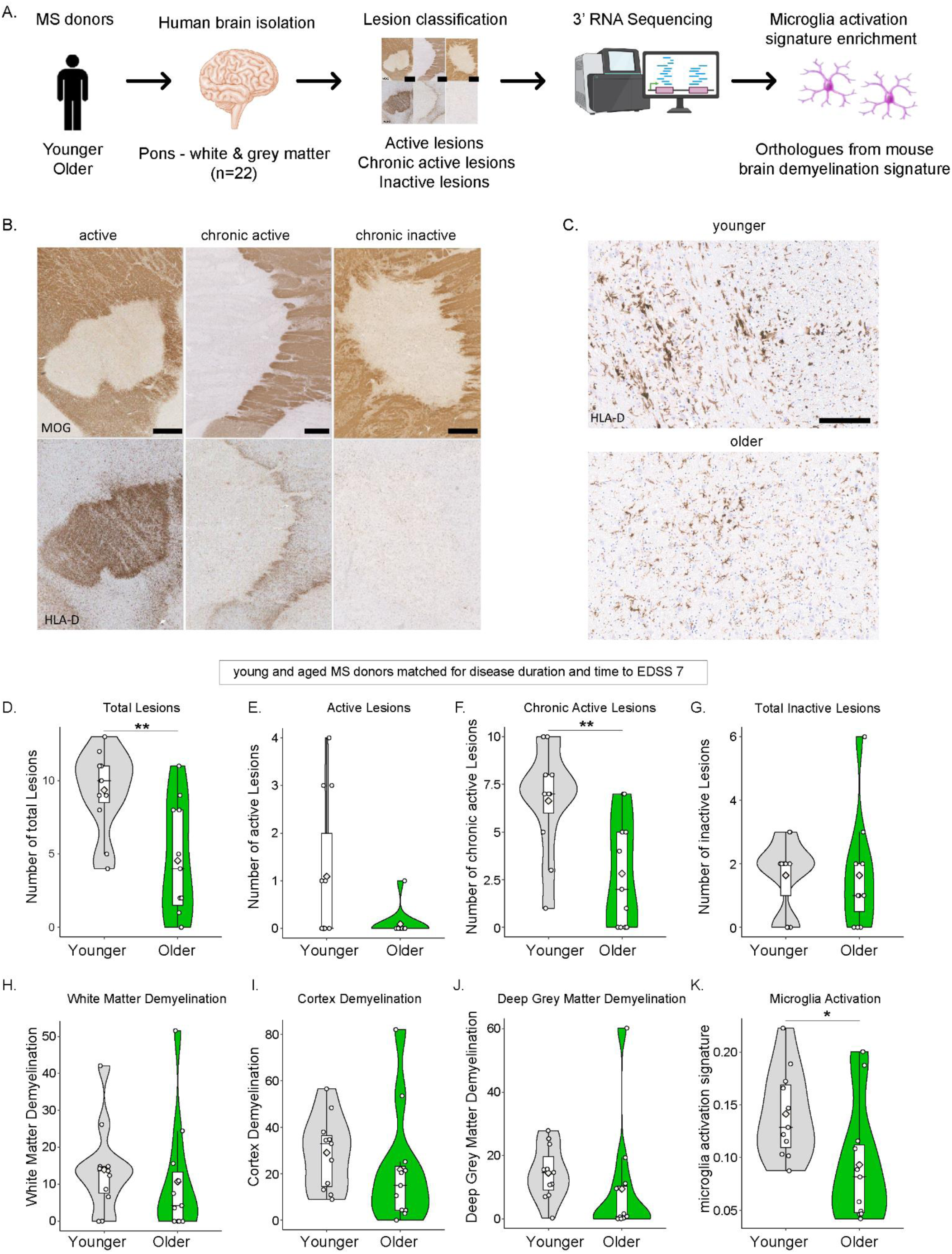
Human MS brain microglia activation is reduced with increasing age **(A)** Experimental design of the human MS RNA-Seq study. Post-mortem brain tissue from younger and older MS donors (n=22) was collected from the pons (white and grey matter). Lesions were classified based on HLA-D immunoreactivity into active, chronic active (including broad-rim lesions), and chronic inactive categories. RNA was extracted and subjected to 3′ RNA sequencing. Microglial activation was quantified using AUCell enrichment of a predefined microglia activation signature derived in this study from the young cuprizone versus young naïve comparison and translated to human orthologues. **(B)** Active, chronic active and chronic inactive demyelinating lesions of the basal pons, characterized by MOG- (myelin) and HLA-D- (immune activation) immunostained sections. Scale bar= 1mm. **(C)** HLA-D- immunostained microglia in younger and older non-lesion MS pons. Scale bar= 200um. **(D–G)** Violin plots showing lesion burden in young and aged MS donors (matched for disease duration and time to wheelchair/EDSS 7). **(D)** Total number of lesions per donor, **(E)** Number of active lesions, **(F)** Number of chronic active lesions and **(G)** Number of chronic inactive lesions, in young and aged MS donors. **(H–K)** Violin plots showing **(H)** White matter demyelination, **(I)** Cortical grey matter demyelination and **(J)** Deep grey matter demyelination, in younger and older MS donors. **(K)** Violin plot showing pontine microglial activation quantified from RNA-seq data using AUCell enrichment of the mouse microglia activation gene signature from brain demyelination RNA-Seq of this study. Each point represents an individual donor. (mean ± SEM, p<0.05, *p<0.01, **p<0.001, ***p<0.0001).

## Discussion

Aging negatively affects myelin, as evidenced by age-related reduction in brain WM volume and decline in structural integrity, contributing to a decline in cognitive functions. Furthermore, WM provides a substrate for immune-mediated diseases like MS, which further burdens spontaneous endogenous CNS repair mechanisms like removal of dead myelin and remyelination by OPCs. Here, we identify microglial aging as a principal factor responsible for age-related loss of beneficial demyelinating and remyelinating brain functions in mice, and provide evidence for reduced microglial activation with increasing age in pons from people with MS. Using in vitro and in vivo demyelination models, we show that aged mice have reduced microglia activation responses to cuprizone demyelination, reduced potential to clear and repair dead myelin and develop a progressive, poorly-demyelinating, non-resolving neuroinflammatory disease in response to sequential episodes of demyelination, instead of alternating cycles of demyelination and remyelination reminiscent of relapsing-remitting MS, as in young mice. We further show that microglia depletion and repopulation using the Csf1r antagonist, BLZ945, was sufficient to recover microglia debris phagocytosis and remyelination in both white and grey matter regions of aged mice and suggest that microglial replacement might represent a promising strategy for the treatment of progressive MS.

Our finding that cuprizone-induced demyelination pathology is significantly delayed in aged mice is consistent with previous studies, in which higher concentrations of cuprizone were used in aged mice to induce pathology with comparable timing and severity as that in young mice^48,49^. We found that demyelination in aged mice was not significantly accelerated by doubling the cuprizone dosage (0.4% w/w), and that OLG damage was efficiently induced by the standard (0.2% w/w) cuprizone dosage, suggesting that CNS rather than peripheral factors are important for the altered demyelination pathology. For purposes of comparison, we therefore used the standard cuprizone protocol in all mice. Delayed demyelination pathology in aged mice could result from higher resistance of mature OLGs to cuprizone-induced apoptosis, or reduced function of aged microglia. In line with a previous report^49^, we found OLG loss to be significant in both young and aged mice, and independent of cuprizone concentration. Furthermore, transcriptomics analysis revealed strong early downregulation of myelin genes in both young and aged brains, while extensive demyelination was only seen in young mice. RNAscope analysis showed robust depletion of PLP-expressing OLGs from early disease in both young and aged corpus callosum. All indications are that OLGs are highly susceptible to cuprizone toxicity in both young and aged mice.

On the other hand, aged microglia showed severe functional defects including impaired microglial activation and resolution responses, myelin phagocytosis and lipid clearance, demyelination efficiency, and failure to facilitate remyelination between sequential demyelination challenges in both cortical GM and corpus callosum WM. Transcriptomics analysis of aged naïve compared to young naïve brains showed upregulated microglial genes including immune mediators, partial DAM signatures and several previously-described age-dependent microglia ADEM genes^56^. This is consistent with previous reports that naturally aged microglia are already activated, “primed”, with an inflammatory profile^57–59^. Age-related microglia activation is observed in humans, and is a feature of normal appearing WM in MS patients^60,61^. However, the activation responses of aged microglia to inflammatory stimulation are weakened, indicating overall reduced plasticity of aged microglia^56,62^. Several reports of increased reactivity of aged microglia to lipopolysaccharide (LPS) and other immune stimuli, measured increased IL-1β and IL-6 expression^63–65^, but this is likely to reflect altered polarization to a damaging inflammatory phenotype rather than increased activation. We previously show that IL-1-polarized microglia represent a damaging phenotype that delays myelin repair under demyelinating conditions, and that inhibition of the TNF-TNF receptor IL-1β signalling axis is sufficient to promote brain repair^23^. Similar macrophage/microglia polarizations were associated with poorly-remyelinating and efficiently-remyelinating lesions, respectively in MS^66^. Indeed, IL-1β and TNF impede lipid catabolism in macrophage-derived foam cells^67^. An imbalanced shift of aged microglia towards a chronically-activated IL-1 phenotype with reduced phagocytosis activity and impaired lipid metabolism, is likely to be a major determinant of the worse outcome of CNS demyelination and repair mechanisms in aged mice, and possibly MS. The evidence also infers that highly efficient removal of damaged myelin by healthy phagocytosing microglia (demyelination), is a pre-requisite for successful remyelination of lesions.

The inability of aged brain to initiate and resolve neuroinflammation or repair of white and grey matter lesions between sequential demyelination episodes in mice is particularly relevant for our understanding of pathology in age-related degenerative diseases such as MS. MS mainly presents clinically as a relapse-remitting disease, in which CNS myelin is a substrate for bouts of peripheral immune cell infiltration. With increasing age, disease clinically transitions to a progressive neurodegenerative form associated with CNS compartmentalized inflammation and failed remyelination^35,54,68^. In MS, evolution to the progressive phase appears to be independent of previous immune system involvement^33,35,68^. Here we show that age-related dysfunction of CNS-resident microglia is a major determinant in, and is sufficient for, driving a progressive disease course in a mouse demyelination model. The relevance of this mechanism for the pathology of MS is substantiated by our finding that disease activity, measured by lesion burden, and microglial activation were markedly reduced with increasing age in the pons of MS donors that had been matched for disease duration and length of time to wheelchair (EDSS 7).

The functional requirement of microglia for remyelination in demyelinating models is previously established^16^. Microglia are self-renewing cells that rapidly repopulate the CNS after depletion by CSF1R inhibition^69^, or genetic knockout^16^. The important therapeutic concept of microglia replacement in experimental disease models^70,71^ and human disease^70^, or innate immune training by a vaccine^72^, has already been introduced. Our finding here that microglia depletion by the CSF1R inhibitor BLZ945 between two episodes of demyelination significantly augmented remyelination in both young and aged mice has important implications for the therapy of MS. Currently approved drugs for MS inhibit immune cells such as T and B lymphocytes of block their trafficking and entry into the CNS, and are effective in early relapsing-remitting stages of MS but show limited effect in slowing disease progression^73,74^. Our results suggest that repopulation or replacement of CNS microglia by cells with full functional capacity, rather than inhibition of established microglia, might be a promising therapeutic approach for removing damaged myelin, controlling inflammation and promoting myelin repair in progressive demyelinating diseases.

---

**Supplementary Figure 1.**
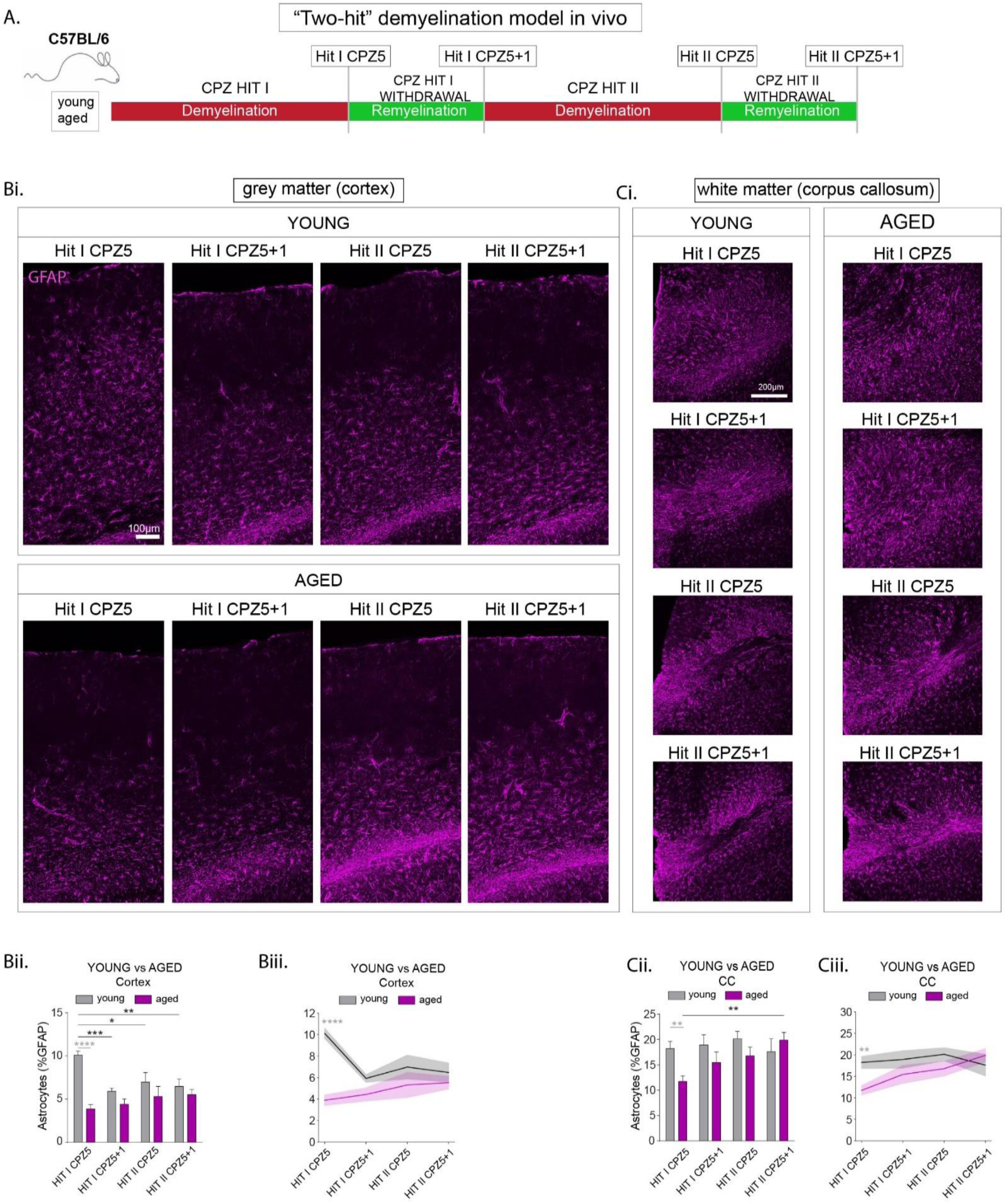
Astrocyte reactivity is delayed and progressive in white and grey matter of aged mice during repeated demyelination episodes **(A)** Two-hit cuprizone demyelination model in vivo. Young (2-month-old) and aged (12-month-old) mice were fed 0.2% cuprizone for 5 weeks (Hit I CPZ5) to induce maximum demyelination, allowed to recover without cuprizone for 1 week (Hit I CPZ5+1), again fed cuprizone for 5 weeks (Hit II CPZ5), and again allowed to recover for 1 week (Hit II CPZ5+1). Confocal image-tiles and analysis of astrocyte reactivity in **(B)** cortical grey matter, and **(C)** midline corpus callosum (CC) white matter, during repeated demyelinating episodes in young and aged mice. **(Bi)** Representative images of GFAP immunostaining of sagittal sections through somatosensory cortex (layers I-VI) (Scale bar: 100μm), and **(Bii–Biii)** quantification of astrocytes (%GFAP immunoreactivity) from longitudinal comparison of young vs aged mice following Hit I (CPZ5, CPZ5+1) and Hit II (CPZ5, CPZ5+1). **(Ci)** Representative images of GFAP immunostaining of coronal sections through midline CC (Scale bar: 200μm), and **(Cii–Ciii)** quantification of astrocytes (%GFAP immunoreactivity) from (longitudinal comparison of young vs aged mice following Hit I (CPZ5, CPZ5+1) and Hit II (CPZ5, CPZ5+1). (n = 4-6 mice/group; mean ± SEM, p<0.05, *p<0.01, **p<0.001, ***p<0.0001).

